# Cellf-Deception: Human microglia clone 3 (HMC3) cells exhibit more astrocyte-like than microglia-like gene expression

**DOI:** 10.1101/2025.08.04.665415

**Authors:** K. K. Rahm, B. S. Kinghorn, M. J. Moody, B. C. Stone, K. C. Strong, B. S. Kim, D. V. Hansen, M. H. Bailey

**Author notes:** First author (K. K. Rahm). Senior author.

## Abstract

Recent advances in Alzheimer’s research suggest that the brain’s immune system plays a critical role in the development and progression of this devastating disease. Microglial cells are vital as immune cells in the brain’s defense system. Human Microglia Clone 3 (HMC3) is a cell line developed as a promising experimental model to understand the role of microglial cells in human diseases including Alzheimer’s and other neurodegenerative diseases. The frequency of HMC3 cell usage has increased in recent years, with the idea that this cell line could serve as a convenient model for human microglial cell functions. Here, we utilize gene-pair ratios from pseudo-bulk and scRNAseq expression data to create predictive models of cell-type origins. Our model reveals that the HMC3 cell line represents various cells, with the highest cell similarity score relating to astrocytes, not microglia. These findings suggest that the HMC3 cell line is not a reliable human microglia model and that extreme caution should be taken when interpreting the results of studies using the HMC3 cell line.

## Introduction

As Alzheimer’s Disease (AD) is a growing global health concern and a leading cause of death in the United States, it is essential to better understand this disease (Weuve et al. 2014; James et al. 2014; “2024 Alzheimer’s Disease Facts and Figures” 2024). AD leads to the breakdown of neurons in the brain and eventually to brain atrophy (Cedres et al. 2020). Recent studies suggest that the brain’s immune system may play a key role in developing AD (Weiner 2025). Microglial cells are known as the first line of defense in the brain’s innate immune system functionality. These cells maintain brain homeostasis and, when working properly, find cells that are diseased or injured (Bohlen et al. 2019; Condello et al. 2015; Vainchtein and Molofsky 2020). Clustering and chronic activation of microglial cells around β-amyloid plaques have long hinted at potential roles for these cells in AD progression (McGeer et al. 1993), and recent identification in genome-wide association studies of many AD risk genes with microglia-specific expression have removed any doubt that microglia are a key cell type that governs AD pathogenesis (Hansen, Hanson, and Sheng 2018). Many genetic and environmental factors can alter the activity and responses of microglial cells, and the mechanisms by which microglia promote or restrain AD development and progression are not fully understood and require further investigation.

AD-relevant microglial activities such as chemotaxis, phagocytosis, lysosome function, and proteostasis can be modeled in vitro using cultured cells. Primary human microglial cells isolated from brain tissue are not commonly utilized since fresh brain tissue is not readily available. Human microglia-like cells known as iMG or iMGL (iPSC-derived microglia-like) cells can be differentiated from induced pluripotent stem cell (iPSC) lines and are increasingly recognized as the best available alternative to primary microglial cells (Wickstead 2023; Abud et al. 2017). However, iPSC maintenance and iMG differentiation are expensive, laborious, and lengthy procedures. Another alternative that allows researchers to produce their (pseudo)cell type of interest in unlimited numbers is to convert primary cultured cells into “immortalized” cell lines with unlimited proliferative potential. Although immortalized cells are not the same as their primary cell progenitors, they may nonetheless serve as useful substitutes that are convenient and inexpensive to culture, and they are more amenable to genetic manipulation to study the functionality of the individual genes.

Currently, multiple immortalized lines of murine microglial cells are available, including the widely used BV-2 and N9 cell lines, both immortalized using retroviral oncogenes (Blasi et al. 1990; Righi et al. 1989; Timmerman, Burm, and Bajramovic 2018). However, interspecific differences between human and mouse immune signaling necessitate the use of human cells. For example, the microglia-expressed AD risk gene and therapeutic target *CD33* has no murine ortholog (Bhattacherjee et al. 2019). Very few immortalized human microglia cell lines have been produced, but one cell line being increasingly utilized by researchers is the Human Microglia Clone 3 (HMC3) cell line (**Figure 1A**).

**Figure 1:**
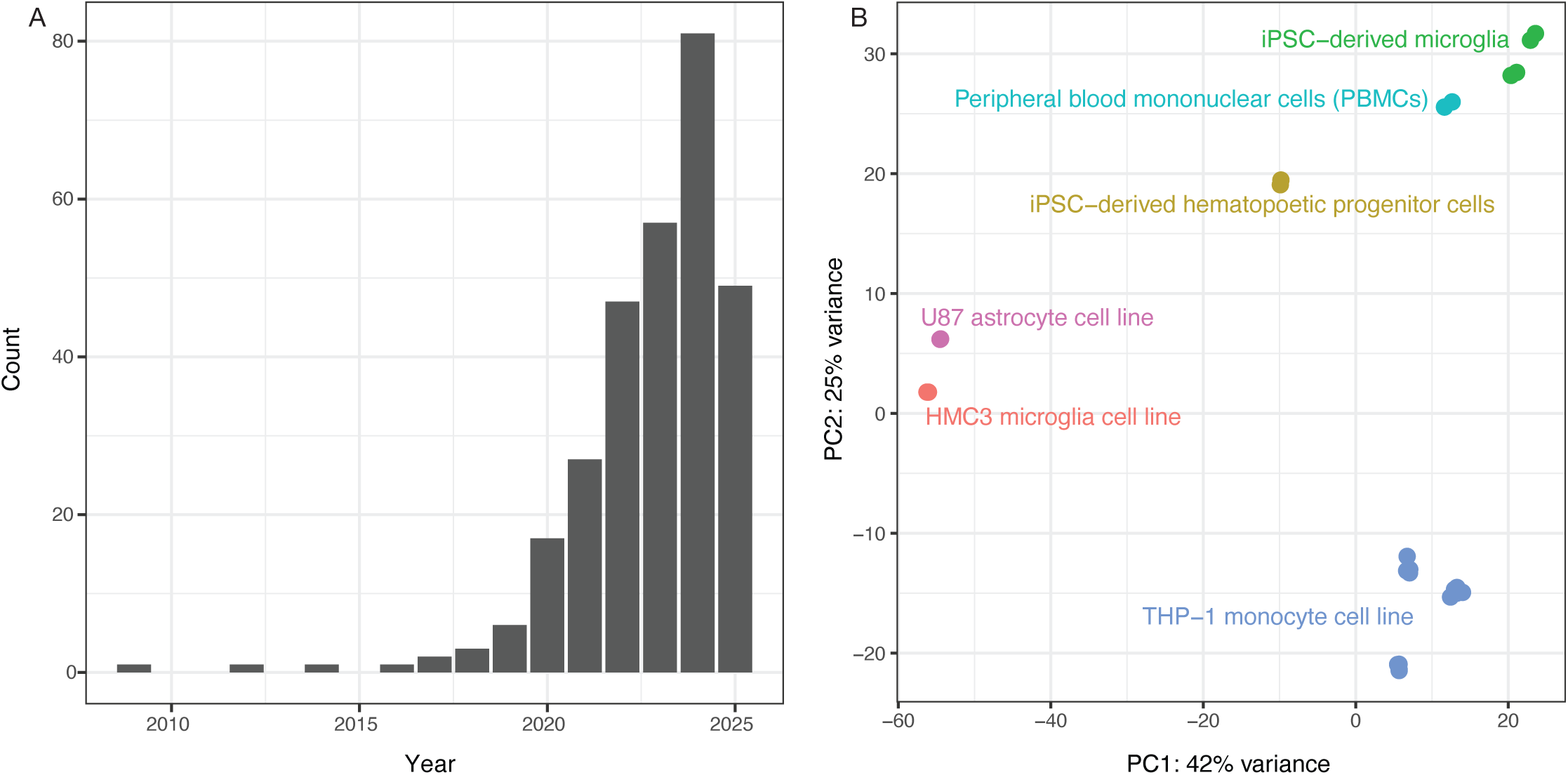
Analysis of previous literature justifies the study. **(A)** PubMed search results for “(HMC3) AND (microglia)” show sharply increasing numbers of publications since 2020, with 57 articles in 2023, 81 in 2024 and 46 so far in 2025 as of July 12. **(B)** Principal components were recalculated using brain cell types from Quiroga, 2022 (GSE181153). We observed an unexpected clustering of the HMC3 microglial cell line with the U-87 astrocyte cell line (red and pink clusters). Note: Adapted from *Synthetic amyloid beta does not induce a robust transcriptional response in innate immune cell culture systems*, I. Y. Quiroga, 2022, http://creativecommons.org/licenses/by/4.0/ (Quiroga et al. 2022).

The HMC3 cell line was created by immortalizing human embryonic microglial cultures with SV40 antigen to further scientific investigation of how microglial cells impact human conditions including Alzheimer’s and other neurodegenerative diseases (Janabi et al. 1995).

Notably, HMC3 cells have also been distributed and used under different names including CHME-3, CHME-5, and C13-NJ cells (Dello Russo et al. 2018). In 2016, it was discovered that ostensible CHME-5 cells being used by several labs at that time were in fact rat-derived cells (Garcia-Mesa 2017), prompting the American Type Culture Collection (ATCC) to authenticate the human origin of the HMC3 cell line (product #CRL-3304).

Motivated by our own interest in using HMC3 cells as potential research tools, we spot-checked some existing HMC3 RNAseq datasets from Gene Expression Omnibus to validate the expression of common myeloid cell markers including *ITGAM* (CD11b), *PTPRC*, *SYK*, *TYROBP*, *CX3CR1*, and Fc receptors. To our surprise, these markers were not expressed, prompting us to examine the research literature for insights as to whether HMC3 cells should be classified as microglia-derived. Indeed, some researchers have noted the lack of expected marker expression in HMC3 cells (Rawat and Spector 2017; Rai et al. 2020), while others reported that HMC3 cells’ transcriptional profile more closely resembled that of U87 cells (Quiroga et al. 2022)—a glioblastoma-derived cell line commonly used to represent astrocyte biology—than profiles of myeloid lineage cells including iMG cells, the monocytic leukemia-derived THP-1 cell line, or primary cultures of microglia, macrophages, or monocytes (see our similar principal component analysis in **Figure 1B**). If the research community increasingly uses the HMC3 cell line to study the role of microglial cells in neurodegenerative diseases, we must present a definitive classification of what type of brain cell it best represents.

The purpose of this paper is to computationally assess the myeloid nature of the HMC3 cell line, specifically distinguishing between microglia and astrocytes or other CNS cell types. Although astrocytes and microglia are physically different and easy to distinguish while examining morphology in vivo (Vainchtein and Molofsky 2020), transformed or immortalized cells cultured in vitro have less distinct morphologies and can have altered traits due to the immortalization process (Kaur and Dufour 2012). are physically different and easy to distinguish while examining morphology in vivo (Vainchtein and Molofsky 2020), transformed or immortalized cells cultured in vitro have less distinct morphologies and can have altered traits due to the immortalization process (Kaur and Dufour 2012). Single-cell and bulk RNA sequencing are powerful tools for distinguishing cell types by measuring transcriptome-wide differences in gene expression levels, providing a robust alternative to morphology-based observations (Haque et al. 2017; Mukamel and Ngai 2019; Ofengeim et al. 2017).

Our strategy for HMC3 cell classification used publicly available RNAseq expression data from multiple studies to determine the proper placement of the HMC3 cell line amongst different lineages of cells within the brain. Microglial cells and other CNS cell types have been sequenced many times for study and classification (Spurgat and Tang 2022), and the resulting datasets are accessible in databases such as Gene Expression Omnibus (GEO) and Genotype-Tissue Expression Portal (GTEx) (Gerrits et al. 2020; Keil, Qalieh, and Kwan 2018).

Our primary goal was to understand if the HMC3 line could be confidently characterized as microglia-derived cells and subsequently used in researching the mechanisms of neurodegenerative diseases such as Alzheimer’s. Given that gene expression directly influences phenotypic features of a cell, including its behavior, we developed two independent Random Forest classifiers to investigate the cellular identity of the HMC3 cell line using gene-ratio comparisons. The first classifier was trained on primary human cells from multiple studies to establish gene-pair rules for distinguishing among cell types and to generate prediction scores reflecting cell-type similarity. When the HMC3 cell line was analyzed with this classifier, it exhibited variability in predicted cell-type scores, with the highest similarity observed for astrocytes. The second classifier was trained on the extensive cohort of cell lines collected by the DepMap project (Tsherniak et al. 2017).

## Materials and Methods

### PubMed trend analysis of “HMC3 AND microglia” publications

We conducted a comprehensive search on the PubMed (pubmed.ncbi.nlm.nih.gov/) research database to identify all articles published between 2009 and July 12, 2025 that included both “HMC3 AND Microglia” as search terms. After downloading the search results, we compiled the number of relevant publications per year and created a table summarizing the annual publication counts.

### Data access and cleaning

RNA sequencing datasets derived from healthy human tissues, a variety of brain cell types, and various cell lines were collected from GEO, DepMap, and several papers (**Supplementary Table 1**). Control and diseased samples from each dataset were used as part of our analysis. All datasets used the HUGO gene symbols, and any datasets that used the human ENSEMBL gene nomenclature were converted through the R IDConverter package to enable gene-pair comparisons among different datasets (Wang et al. 2021). Additionally, we restricted genes within the datasets to those where 80% or higher of the samples had a non-zero observation in the specified gene. This restricted our training datasets to 8,723 genes.

### Principal Component Analysis

**Figure 1B** in this manuscript mimics Figure 1 from Quiroga et al. (2022) (GSE181153) but with fewer cell type comparisons (Quiroga et al. 2022). First, the gene-counts table was converted to transcripts per million. As performed by Quiroga et al., only genes with greater than 100 transcripts per million were kept in the analysis. Samples were then restricted to only include control non-treated samples, i.e., only “NONE_NONE” labels were kept. Additionally, in order to recreate the Quiroga PCA, iPSCs were similarly not included in the plot, despite the raw data including iPSCs. Of note, when the iPSC lines were included, they cluster with U87a and HMC3 lines (data not shown). Once samples and genes were removed, gene count data were processed using DEseq2 (Love, Huber, and Anders 2014), followed by variant stabilization (“vst()” function). Following the variant stabilization step, the ‘plotPCA()’, a native function in DEseq2, was used to generate the Principal Component plots.

### The primary-cell Random Forest classifier

A primary cell Random Forest classifier was developed using the multiclassPairs R package (v0.4.3), which implements a rule-based classification framework with gene-pair comparisons (Marzouka and Eriksson 2021). The classifier was built from five publicly available datasets, comprising a total of 258 healthy and diseased human samples. From the Galatro et al. dataset (GSE99074), 65 microglia samples were included. The Gosselin et al. dataset (phs001373.v1.p1.) contributed 46 microglia and 13 monocyte samples. The Srinivasan et al. dataset (GSE125050) provided 19 adult astrocytes, 27 endothelial, 25 microglia, and 42 neuron samples. From the Zhang et al. dataset (GSE73721), 9 adult astrocyte, 4 microglia, and 1 neuron sample were included. Lastly, the Costa-Verdera et al. dataset (GSE253820) contributed 4 adult astrocyte and 3 neuron samples (**Supplementary Table 1**). After preprocessing, each sample was annotated with its respective cell type and dataset of origin. All data were merged into a single gene expression matrix, with genes as rows and samples as columns, for input into the multiclassPairs workflow.

The ReadData function from the multiclassPairs R package was implemented to structure the combined matrix into an appropriate format with correct labeling for downstream analysis. The data was randomly partitioned into training (60%) and testing (40%) sets, ensuring no sample overlap between sets.

To reduce dimensionality and improve classifier performance, feature selection was performed using the ‘sort_genes_RF’ function. This function ranks genes based on their importance in differentiating cell types, employing a Random Forest-based feature ranking strategy of importance scores computed by the embedded ranger package. A total of 2000 trees were used in the feature selection step. The top-ranked genes were selected for further analysis using the summary_genes_RF function, which indicated that 85 genes from the “altogether” category and 100 genes from the “one-vs-rest” category provided optimal rule coverage for downstream training.

From these genes, binary classification rules were generated by forming gene pairs, where a rule was defined as “Gene A < Gene B.” These rules were sorted based on their discriminative power using the sort_rules_RF function. To ensure robustness, rule selection was performed using both one-vs-rest and all-vs-all approaches, allowing for differential weights of all cell types to reduce bias due to sample imbalances.

The final classifier was then trained using the “train_RF” function, with model parameters optimized through the “optimize_RF” function. Specifically, the model was then constructed of 1,000 trees, with a gene repetition limit of one to ensure rule diversity. A total of 100 rules derived from the one-vs-rest scheme and 85 rules from the altogether scheme were selected for training. Boruta-based feature selection was enabled to exclude non-informative rules (Kursa, Jankowski, and Rudnicki 2010). Additionally, probability estimation was activated, allowing the model to output class scores instead of categorical predictions. The resulting model was composed of 452 binary rules (gene-pairs) across all classes (**Supplementary Figure 1**).

### Primary cell-line RF classifier

Concurrently, we generated a cell-line-based Random Forest classifier as before, but restricted the training set to cell lines. Specifically, we used data from DepMap (Tsherniak et al. 2017). Built on the foundation of the Cancer Cell Line Encyclopedia (Cancer Cell Line Encyclopedia Consortium and Genomics of Drug Sensitivity in Cancer Consortium 2015), the DepMap project’s goal is to model and explore the molecular dependencies that govern cell immortality and shed light on which genes are necessary and sufficient to keep cancer growing forever so that they may be exploited for future treatments. For this project, we used 174 cell lines collected from the DepMap Download portal at https://depmap.org/portal/data_page/?tab=currentRelease version 24Q4 under the expression tab. We downloaded and integrated data from “Model.csv” (the collection of metadata used to describe the cell types and their origins, including sex and tissue of origin) and “OmicsExpressionProteinCodingGenesTPMLogp1.csv” to get the expression data. Gene numbers were removed, and genes were subset as above. Additionally, we subset our data to cell types that originated in the brain/central nervous system, or blood cancers in order to classify cells as neural or myeloid in nature. 22 gene names were missing from the DepMap data and were not considered in the training: “RBM14_RBM4”, “FPGT_TNNI3K”, “BCL2L2_PABPN1”, “TEN1_CDK3”, “PPT2_EGFL8”, “RTEL1_TNFRSF6B”, “SENP3_EIF4A1”, “P2RX5_TAX1BP3”, “STX16_NPEPL1”, “DLEU1”, “ERV3_1”, “HLA_A”, “HLA_C”, “HLA_DMA”, “HLA_DQB1”, “HLA_DRB1”, “HLA_F”, “CHKB_CPT1B”, “ST20”, “ANKHD1_EIF4EBP3”, “ZNF286B”, and “JMJD7_PLA2G4B”. This left 8702 genes for training the tissue of origin data.

Note “ACH-000075” or the U87 cell line *is* part of the DepMap dataset. We ensured that “ACH-000075” was not included in the training of the DepMap model, but was confined to the test dataset. This also provides some justification for not performing cross-validation on our data, where U87 might eventually be included to optimize the model.

After following the same methodology above, 41 rules from 82 gene-pairs were used to build our classifier (**Supplementary Figure 2**).

### Model testing and evaluation

Upon the creation of two classifiers, a primary cell classifier and cell-line classifier, we tested RNAseq data generated from many different studies (Gupta et al. 2024; Schirmer et al. 2019; Masuda et al. 2019; Quiroga et al. 2022; Armanville et al. 2025; Baek et al. 2021, 2022; Abud et al. 2017) to evaluate the performance of our model and assess its validity. The Caret::confusionMatrix function was utilized to generate model performance metrics for both the training and test sets (**Supplementary Tables 2 and 3**).

### Testing for rat sequencing reads

FASTQ files were downloaded to a Google bucket from the Sequence Read Archive (SRA) for all HMC3 samples tested (GSE181153, GSE275256, GSE155408, specifically, SRR12347826, SRR12347827, SRR12347828, SRR15301012, SRR15301013, SRR15301082, SRR15301083, SRR30311414, SRR30311415, SRR30311416, SRR30311417, SRR30311418, SRR30311419, SRR30311420, SRR30311421, and SRR30311422). To test whether samples had more RNA sequencing reads that aligned to Rat (Rattus norvegicus) or Human (Homo sapiens) reads, we used xengsort (Zentgraf and Rahmann 2021) (https://gitlab.com/genomeinformatics/xengsort) to characterize reads according to their alignment preference. We used Ensembl GRCh38 release 114 cDNA and DNA FASTA files for the human reference and GRCr8 release 114 cDNA and DNA sequences for the rat genome reference. Briefly, the xengsort pipeline uses a memory-intensive step to index the reference genomes, and then uses Cuckoo hashing for rapid assessment of k-mer fidelity in order to classify raw reads to the different references for each sample.

### Code availability

The code used to clean and analyze these data is available at https://github.com/MHBailey/Cellf_deception.

## Results

### Data collection

Data was collected from fourteen different studies involving brain cells and cell lines (Srinivasan et al. 2020; Galatro et al. 2017; Zhang et al. 2016; Gosselin et al. 2017; Costa-Verdera et al. 2025; Tsherniak et al. 2017; Gupta et al. 2024; Schirmer et al. 2019; Masuda et al. 2019; Quiroga et al. 2022; Armanville et al. 2025; Baek et al. 2021, 2022; Abud et al. 2017) (**Supplementary Table 1**). In total, 432 samples from six of these studies were used to train two different classifiers–a classifier for cells collected from human tissues and another classifier from cell lines. (**Materials and Methods**).

### Building a primary tissue-type Random Forest classifier

A statistical R package called multiclassPairs was used to build these classifiers (Marzouka and Eriksson 2021) (**Materials and Methods**). Briefly, multiclassPairs builds Random Forest classification models based on the relationship between two genes instead of single gene quantities. Gene-pair ratios are less subject to batch effects (Ellrott et al. 2025) and thus more comparable when including many studies. Specifically, we leveraged both bulk-sorted and single-cell RNAseq (scRNAseq) data from multiple studies to create our classifiers.

#### Primary tissue classifier training

258 samples were derived from tissue material (**Supplementary Figure 1**). Using RNAseq information from these samples (bulk and single-cell transcriptomics), we built a Random Forest using multiclassPairs. We used a 60:40 training-to-test ratio. In the training data, only one labeled astrocyte sample was predicted as an endothelial cell, and one neuron was misclassified to be microglial in origin (**Supplementary Table 2**). The testing confusion matrix showed promising data with an overall accuracy of 98.08% (CI 0.9323-0.9977) and high balanced accuracies per cell type (>95.83% for each cell type), showing that the majority of samples were classified as their correct cell type.

The primary classifier used 452 gene-pair rules to classify labeled cell types (Figure 2A). Using these gene-pair rules, most cell types have distinguishable groups of gene rules that lead to their prediction; notably, gene-pairs-features are less sensitive to the data platform (Figure 2B). Overall balanced accuracy, sensitivity, and specificity provided strong evidence that our model could correctly classify the cell-type of origin for data derived from specific cells (Figure 2C**, Supplementary Table 2**). For our purposes, we wished to use this classifier as a means to determine the fidelity of established microglial cell lines to the human-derived cellular material.

**Figure 2:**
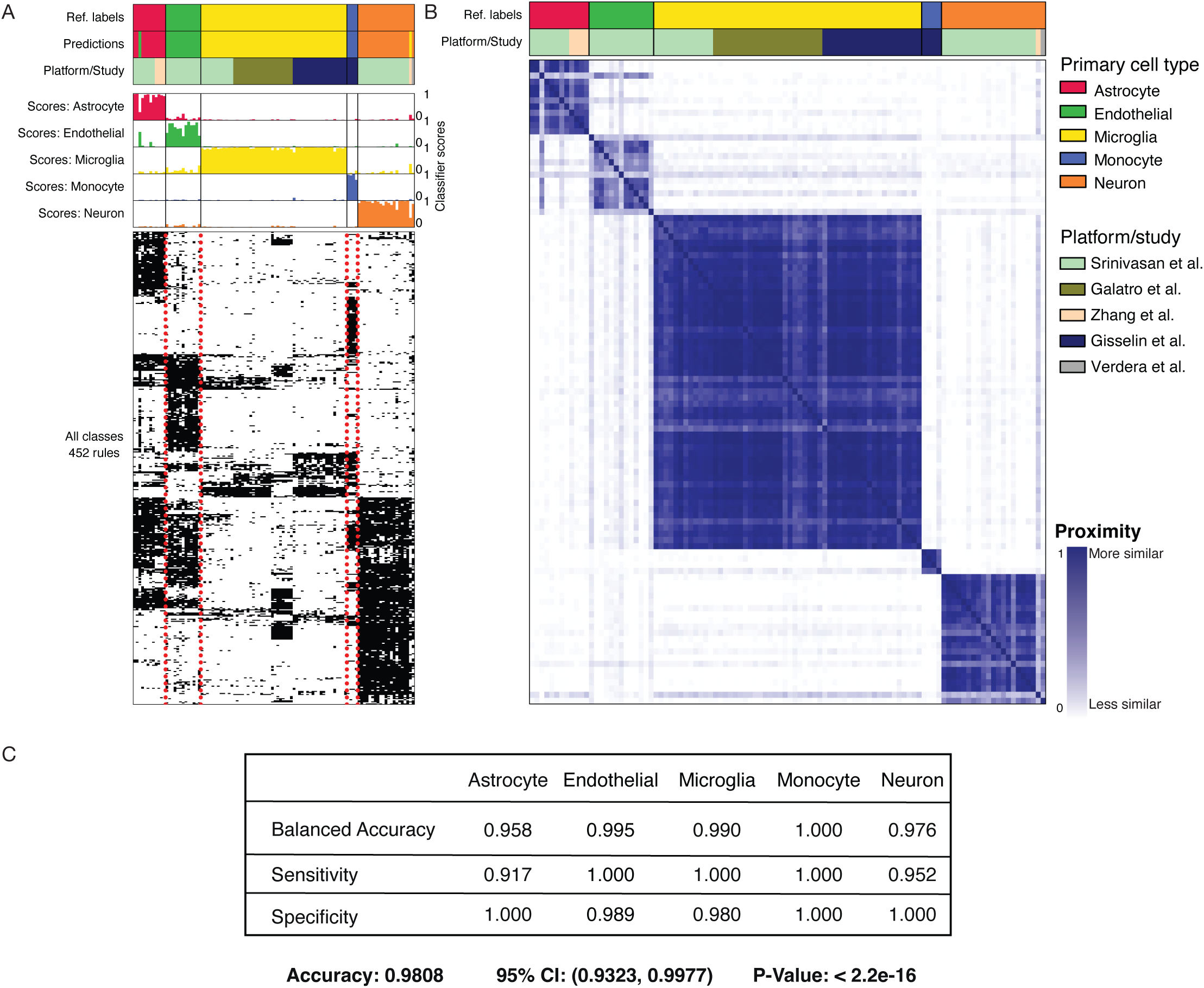
multiclassPairs cell-type classification on the test dataset from primary tissue. **(A)** Three subpanels display the results of the classifier on cells derived and/or sorted from primary tissue. The top panel contains three rows: “Ref. labels” display the cell type assigned by each respective study, “Predictions” display the calls of our classifiers on a set-aside test set, and “Platform/Study” displays how different studies span different cell types. The middle panel displays each sample’s rule activation score (columns) or classification score. The bottom panel displays the binary heatmap of 452 gene-pair decision rules. **(B)** A proximity matrix heatmap portrays similarity clusters of the RF classifier using out-of-bag samples. The top, “Ref. labels” display cell type assigned by each respective study, and the “Platform/Study” row displays how different studies span different cell types. The heatmap shows the cell types mapped onto themselves, and each sample is given a proximity value with every other sample. **(C)** A confusion matrix displaying the specificity, sensitivity, and balanced accuracy of the classifier for each cell type, along with the overall accuracy, confidence interval, and p-value. A more complete table is found in **Supplementary Table 2**.

#### Validate Primary Tissue Classifier with additional datasets

To further assess the validity and accuracy of the RF classifiers, we tested data from two unrelated studies as independent test sets (Figure 3). The first dataset from Masuda et. al contains gene expression data for seven samples of human microglia cells from both healthy and diseased patients (Masuda et al. 2019). The second dataset from Schirmer et al. contains gene expression data for nine samples of human astrocyte cells from healthy patients (Schirmer et al. 2019). Despite higher imputation levels (291 missing genes for Schirmer et al. and 292 missing genes for Masuda et al., i.e., more than half of the rules missing, see **Materials and Methods**), all samples from both studies were correctly identified as their known cell type of origin (Figure 3). In the Masuda et al. study, the microglial cell prediction score for each sample was above 95% (Figure 3A). These consistent results for every sample demonstrate the high efficiency of the RF classifier to make cell-type-specific predictions even with missing data. Similarly, 8 of 9 astrocyte samples from the Schirmer et al. study were correctly classified by a cell prediction score of at least 0.881 (Schirmer et al. 2019). The highest astrocyte cell prediction score was 0.957 (Figure 3B). Interestingly, samples with the lowest predictive score, C7 Astrocyte, C9 Astrocyte, C2 Astrocyte, and C8 Astrocyte, had the fewest astrocyte-labeled cells from their single-cell experiments (average number of labels astrocytes were 31.5 cells compared to 289 cells in the rest of the samples: C4, C5, C1, C6, C3. Averages generated from Schirmer et al.’s Supplementary table 4. These lower cell counts may have led to their less consistent predictions. Overall, these two validation sets provide strong evidence for the strength of our classification tool.

**Figure 3:**
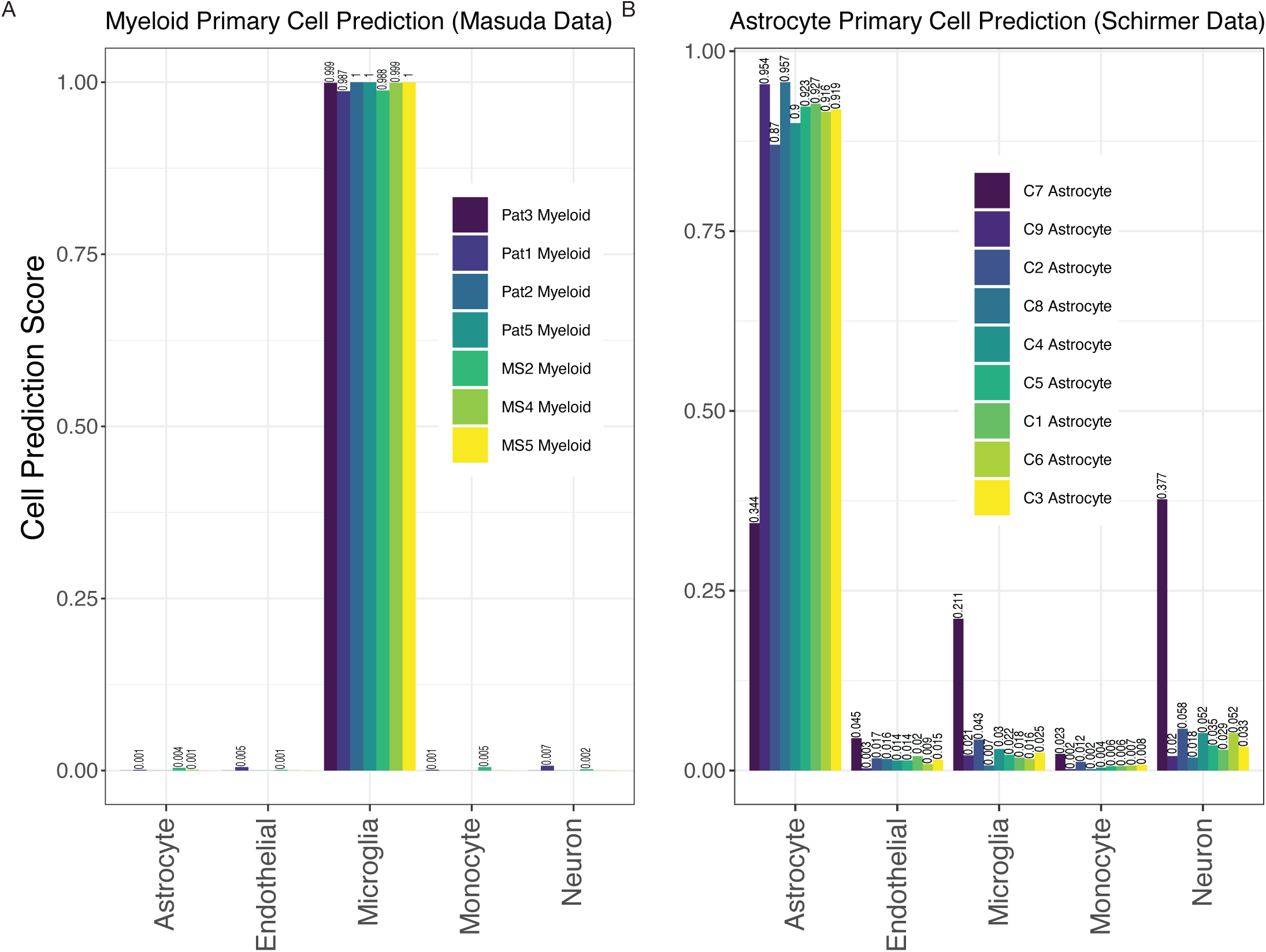
multiclassPairs cell-type classification on independent validation sets of primary tissue. **(A)** Seven human microglia samples from Masuda et al. were run through the classifier. Cell prediction scores for microglia range from 0.95 to 0.989. **(B)** Nine human astrocyte samples from Schirmer et al. were run through the classifier. Cell prediction scores for astrocytes range from 0.334 to 0.957. Values closer to 1 indicate stronger cell assignment congruity.

### Classification of the HMC3 Cell Line

We next employed our primary cell-based RF classifier on HMC3 cell line expression datasets to determine how our model classifies this line. We began with four HMC3 samples from GSE181153 by Quiroga et al. (2022). Our model’s predictions for HMC3 cells based on gene-pair expression patterns were less conclusive than for our test cases of primary cell isolates, but the highest cell prediction scores suggested that HMC3 cells were astrocyte-derived, with microglial derivation receiving the second highest scores (Figure 4A). The cell prediction scores for HMC3 cells representing microglia-derived cells ranged from 0.21 to 0.22 while the range of scores for HMC3 cells representing astrocyte-derived cells was 0.45 to 0.46 (Figure 4A). Thus, we can see that the HMC3 cell line exhibits a heterogenous profile, and it may be most appropriately classified as an astrocyte-derived line according to its gene expression profile. It is known that the immortalization process can cause significant genetic changes and therefore alter behavioral characteristics of immortalized cells (Voloshin et al. 2023). When we utilized the primary cell-based RF classifier to analyze iMG samples from the same study (Quiroga et al. 2022) (withheld from the original training data), iMG cells were confidently classified as microglial, with predictions ranging from 0.884 to 0.918 (Figure 4B). It is evident that iMG cells, though tedious to produce, provide a much more accurate experimental model of microglia than the HMC3 cell line. Notably, the HMC3 cell line, while previously considered a microglia model, does not effectively capture microglial characteristics and instead presents a transcriptionally diverse profile more closely aligning with astrocytes.

**Figure 4:**
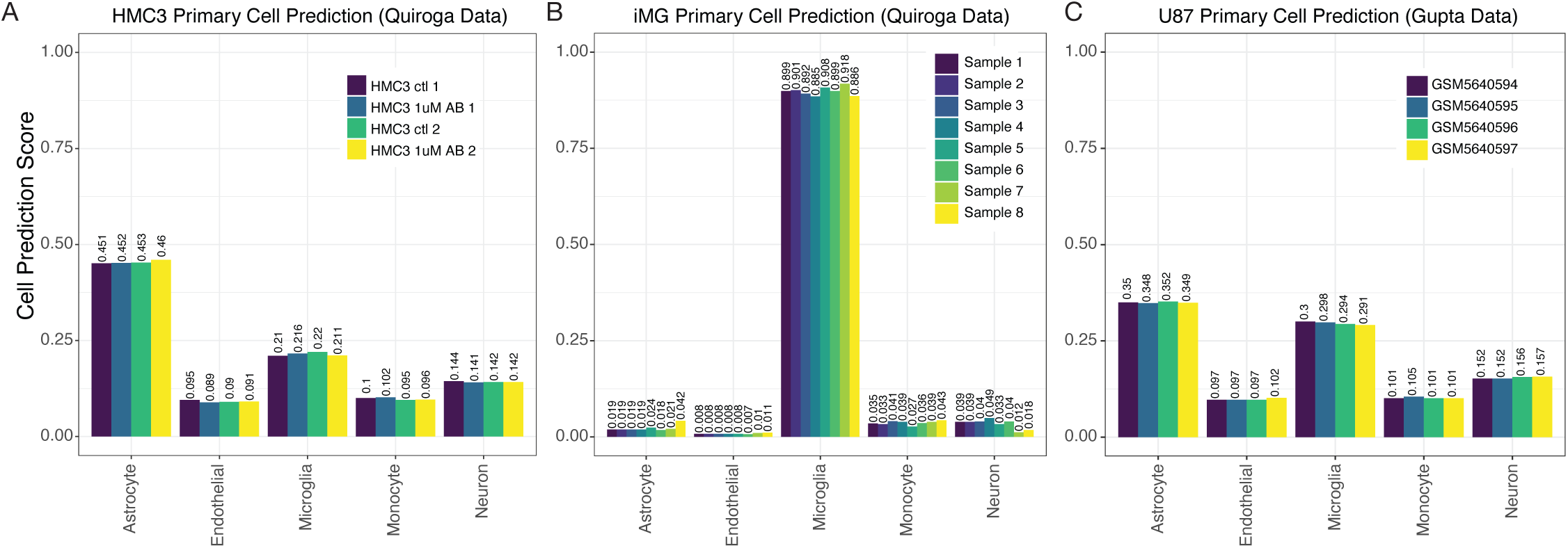
multiclassPairs cell-type classification identifies human microglia cells. **(A)** A dataset of the HMC3 cell lines generated by Quiroga et al. was run through the classifier. A barplot presents an assorted composition of cell prediction scores, identifying most closely with astrocytes. **(B)** Again, an expression dataset of iMG cells was classified by our Random Forest prediction, and rule activation scores are presented in a boxplot. **(C)** The same scores are presented for the classification of U87 cells generated by Gupta et al.

To further explore the original finding from Quiroga et al. that HMC3 cell profiles clustered more closely with U87 glioblastoma-derived cells than with myeloid cells, we also tested U87 cell line profiles in our primary cell-based RF classifier. The U87 samples were also given an ambiguous classification by our tool, with astrocyte scores at ∼0.35 and microglia scores at 0.29–0.3. Interpreting these results, we hypothesized scenarios that might explain the data. First, the immortalization process could put U87 and HMC3 cell lines onto a similar transcriptomic trajectory. This could be supported by our PCA analysis when including iPSCs, which showed iPSCs clustering with U87 and HMC3 cells (data not shown). Second, the original Quiroga PCA analysis could have resulted from a laboratory error of the mixing of U87 and HMC3 cells.

To address the latter possibility, we extended our analysis to two additional datasets that sequenced HMC3 cell lines and deposited their data (Armanville et al. 2025; Baek et al. 2021, 2022). Similar to the HMC3 samples from Quiroga et al., HMC3 samples from these two datasets suggested that HMC3 cells reflect astrocyte expression more than microglial expression, with predictive scores for astrocyte derivation again being ∼2-fold higher than for microglial derivation (Figure 5A **and 5B**). In fact, predictive scores for neuron derivation were as high as the scores for microglial derivation in these HMC3 samples.

**Figure 5:**
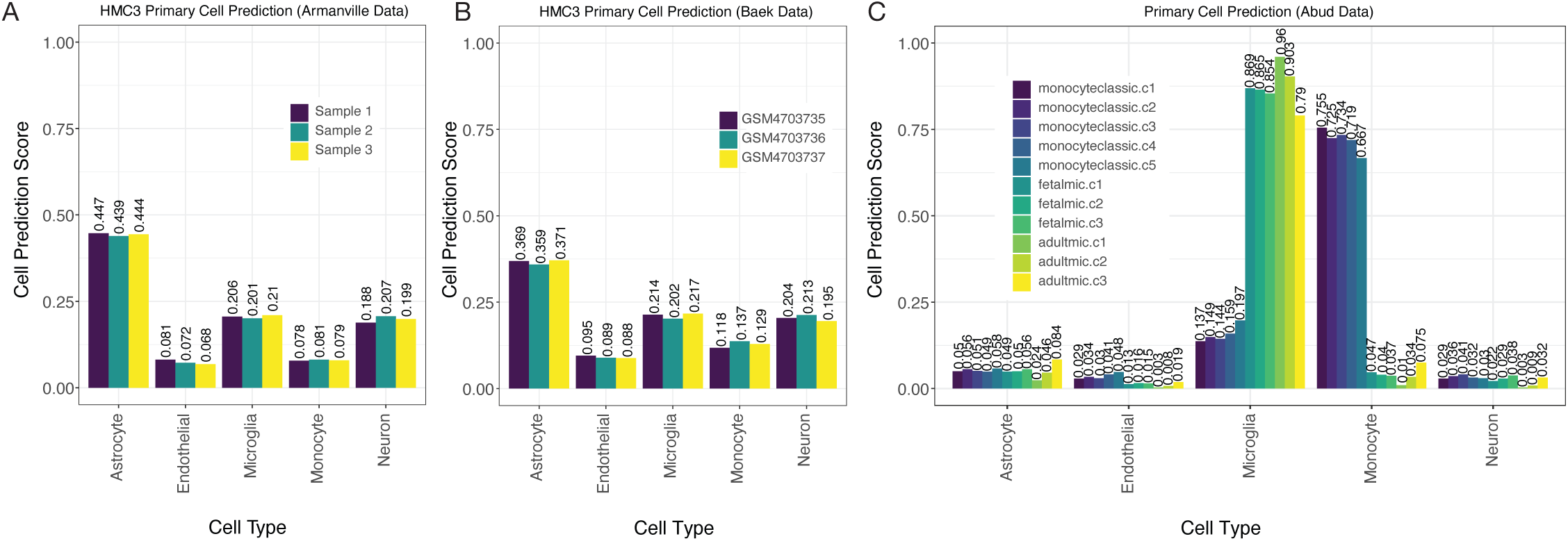
multiclassPairs cell-type classification identifies human microglia cells (A & B). Two additional datasets of the HMC3 cell lines run through the classifier present an assorted composition of cell prediction scores, and identify most closely with astrocytes. **(C)** An expression dataset of monocytes and microglia (fetal and adult) cells was collected from human tissue and analyzed by our primary cell classifier, which shows clear distinctions between microglia from monocytic cells. Sample names provided by the manuscript, and indicators in the sample name indicate the origin of the cells collected: 5 classic monocytes, 3 fetal <u>mic</u>roglial samples, and 3 adult <u>mic</u>roglial collections.

To further explore the specificity of our model in accurately classifying both fetal and adult microglial cells, we incorporated an additional test using the Abud et al. dataset (2017). Their study included gene expression profiles of monocytes as well as induced microglial (iMG) lines derived from induced pluripotent stem cells. Abud et al. demonstrated that these iMGs closely resembled primary human microglia in their transcriptional profiles (Abud et al. 2017). Due to the thoroughness of their sequencing across many facets, we were able to utilize their data to represent both fetal and adult microglial populations in our classifier (Figure 5C). Upon analysis, our classifier successfully distinguished between monocyte samples and microglial lines, in addition to accurately classifying both fetal and adult microglial cells (Figure 5C). These findings further support the robustness of our gene-pair classifier across data platforms and its relative immunity to batch effects while also confirming its ability to accurately identify microglia regardless of developmental stage.

### Classification of a cell-type classifier using DepMap cell lines

To address the possibility that HMC3 cells and U87 cells clustered together mainly because they are transformed/immortalized cell lines and not because they have similar cell origins, we built another RF classifier using only transformed cell lines derived from human cancers of myeloid cell lineages or of the brain (neural, astroglial, or oligodendroglial cancers). To build our transformed cell line-based RF classifier, we used cell lines studied in the DepMap cell line project (Tsherniak et al. 2017). Similar to above, we used multi-class pairs to build a gene-pair classifier, allowing us to compare the DepMap sample to the datasets we collected above (**Materials and Methods**). Of the 174 different cell lines from CNS/Brain or Myeloid in the DepMap dataset, we used 56 CNS/Brain labeled cell lines and 48 Myeloid cell lines to train the gene-pair Random Forest classifier (**Supplementary Figure 2A and 2B**). The test set revealed an F1 statistic of 0.978, an accuracy of 97.1%, a sensitivity score of 1.00, and a specificity of 0.923 when using the remaining cells as a test set (**Supplementary Table 3**). Of note, the two cell lines that were misclassified in the test set were the HAP1 line (ACH-002475) and HDMYZ (ACH-000190, **Supplementary Figure 2C and 2D**). Both lines have lineages labeled by DepMap to be Myeloid in nature, but were better classified with the Brain/CNS labels in our classifier. Interestingly, HAP1 is a cell line derived from the chronic myeloid leukemia line, KBM-7 (Kotecki, Reddy, and Cochran 1999), and is a near-haploid cell line, making it easier to introduce mutations into a model system. Interestingly, the HAP1 arose from the failed attempt to induce pluripotency in KBM-7 by the overexpression of OCT4, SOX2, MYC, and KLF4. However, the resulting cell lost its hematopoietic markers (Carette et al. 2011), which agrees with our misclassification. The HD-MY-Z cell line, originally derived from a pleural effusion taken from a 29-year-old Hodgkin’s patient (Bargou et al. 1993), has also undergone scrutiny, and is now considered to be more AML in origin (Drexler et al. 2018). Drexler et al.’s commentary suggests these cells are more typical of myelomonocytic cells. Our classifier suggests a different origin that warrants further investigation. These results suggest that our classifier can accurately distinguish between cells and can capture known discrepancies in the field.

When we tested HMC3 lines datasets (none of which are in the DepMap data), we observed close alignment with CNS/Brain cancer-derived cell line profiles, a result that is counter to the non-CNS origin of microglial cells (Figure 6A**, 6D, and 6E**). Similarly, we also observed that U87 cells were predicted to originate from a CNS/Brain lineage, as expected (Figure 6B). U87 lines are part of the DepMap project but were withheld from our original training data to be used as a test case. The iMG cell profiles reported by Quiroga et al. or by Abud et al. aligned much more strongly with myeloid lineage cancers than CNS/Brain cancer cell lines, a result in line with expectation (Figure 6C **and 6F**). Finally, from tissue collected by Abud et al., we show that monocytes, fetal microglia, and adult microglia all classify with myeloid cell line profiles, according to their natural origins (Figure 6F). Overall, the transformed cell line-based classifier provides strong additional evidence that HMC3 cells lack the microglial-like identity or transcriptomic signatures sufficient to warrant their classification as a useful cell line for microglial research.

**Figure 6:**
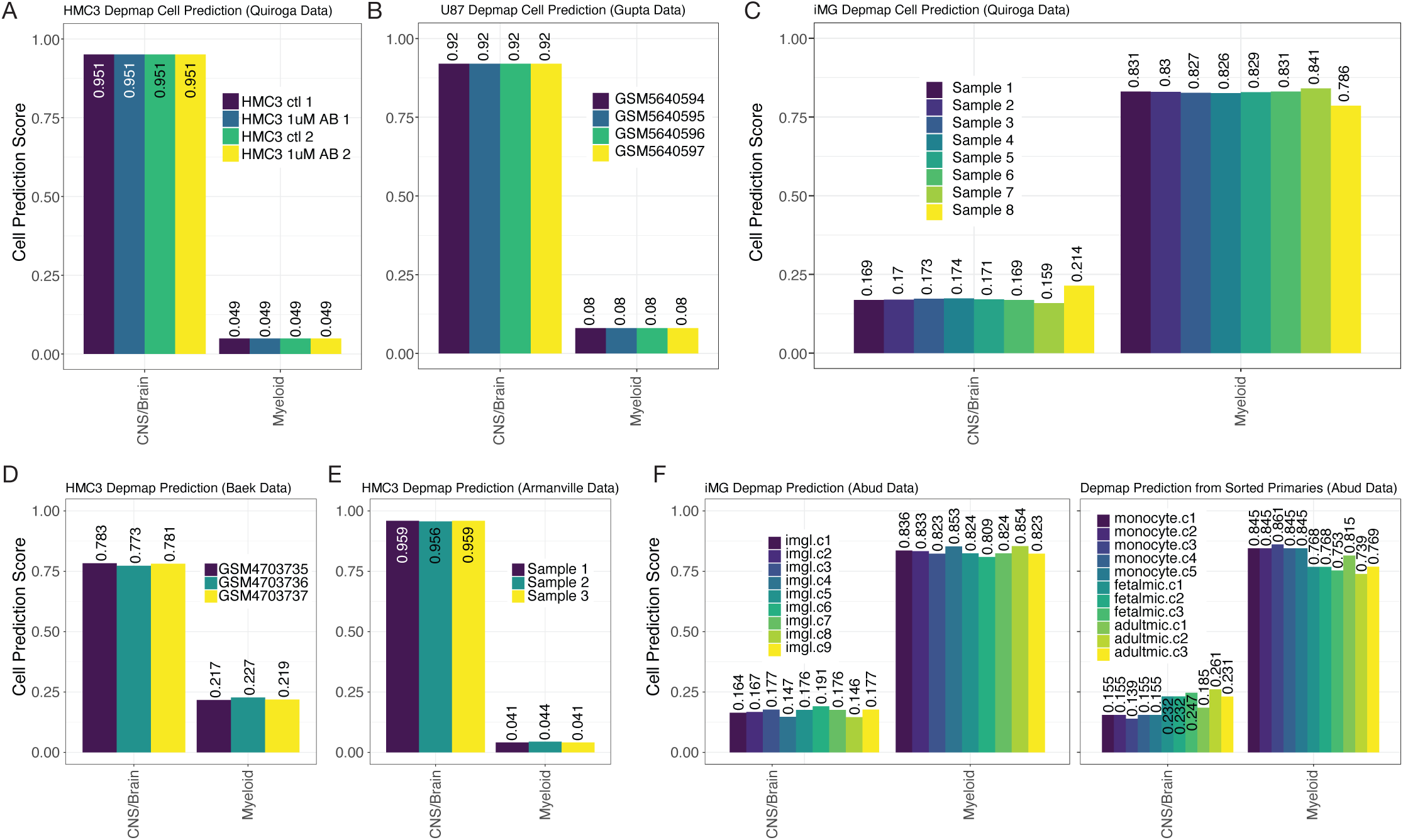
DepMap cell of origin classifier distinguishes myeloid and CNS/Brain cell lines. 7 panels from 5 studies show how the DepMap classifier assigns predictions to many different data types in the exact same manner as Figures 3-5. **(A)** HMC3 lines from Quiroga et al., **(B)** U87s from Gupta et al., **(C)** iMG (induced microglial lines) from Quiroga et al., **(D)** HMC3 lines from Chai et al., **(E)** HMC3 lines from Armanville et al., and (**F)** iMG (induced microglial line), monocytes, fetal microglia, and adult microglia samples from Abud et al.

### Classification of reads in HMC3 cell lines

Finally, one important aspect that needs to be addressed is the possibility that all three HMC3 datasets analyzed in this manuscript were actually derived from rat glioma-derived cells, a widespread problem previously described for some neural cell lines (Garcia-Mesa et al. 2017). Plenty of sequencing reads from rat cells will align fine with the human genome. While ATCC consistently provides high-quality cell lines for labs, we performed a quick assessment of RNA read quality and checked alignments for quality control purposes. Because of the previous challenges with deriving cell lines from Rattus norvegicus (rat), we assessed read origins based on the k-mer fidelity. To accomplish this task, we used a well-designed tool called xengsort (Zentgraf and Rahmann 2021). Briefly, xengsort uses a large alignment-free k-mer store, i.e., a large key-value hash table, to assess whether the DNA or RNA reads best align to one, both, or neither of two references provided. Because of historical problems, it must be addressed explicitly by aligning the HMC3 sequencing reads with human vs rat genomes and analyzing mismatches to ensure that the HMC3 datasets are in fact, human-derived cells and not rat-derived cells. Results from this analysis clearly show that data from the Armenville et al. and Quiroga et al. studies are primarily from the Homo sapiens genome (mean of 98.53% and 98.07%, respectively, Figure 7). Surprisingly, the Baek et al. study did show many shared k-mers with the Rat reference (mean 30.69% shared between human and rat); but only 0.13% of the k-mers were rat-specific. This may be confirmed in our classifier since the Baek et al. samples showed weaker predictions (Figure 6D).

**Figure 7:**
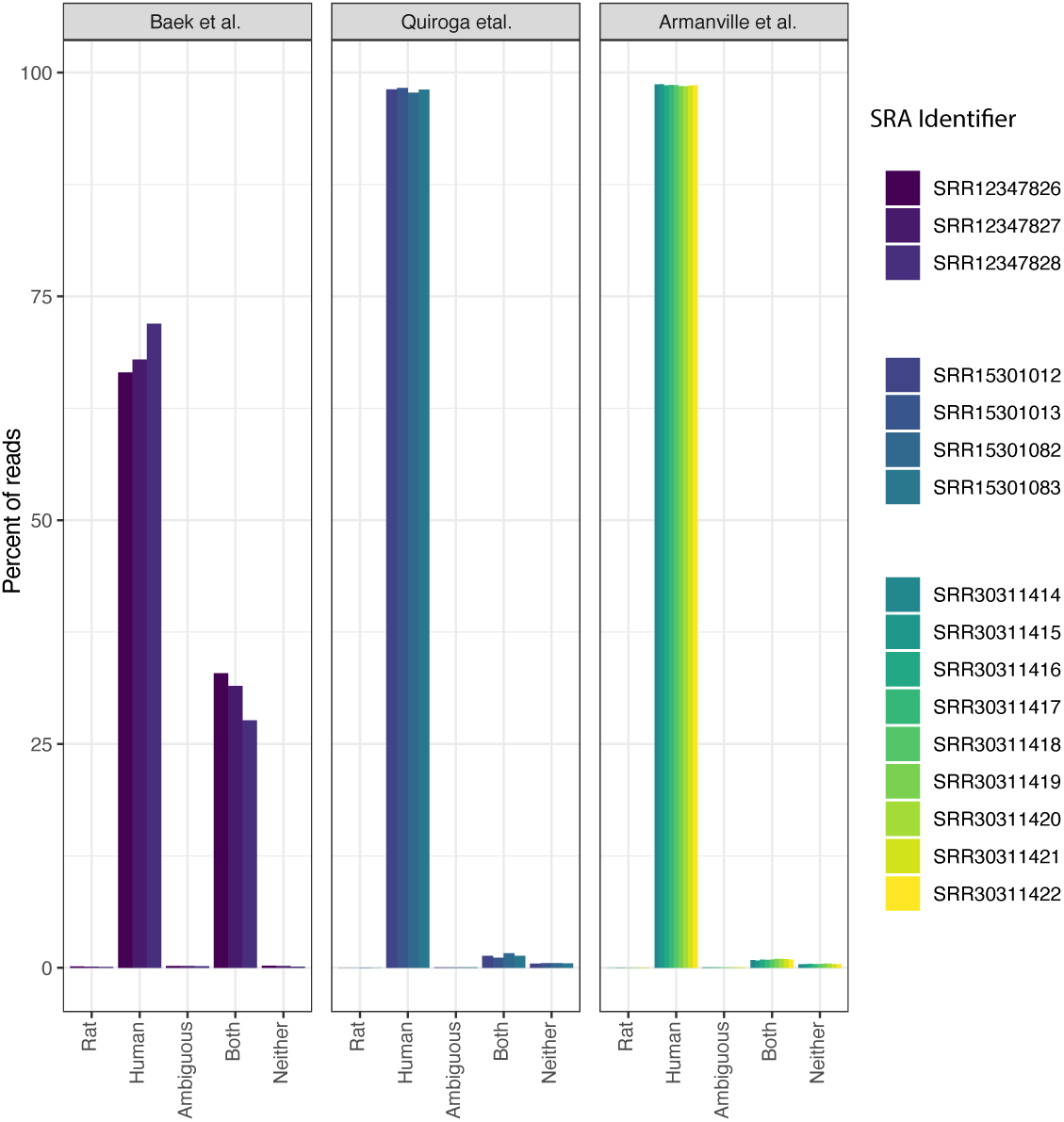
Sequence read quality of the HMC3 cell lines. Bar charts for the three HMC3 studies show the percentage of RNA sequencing kmers that match with the Rat transcriptomic reference (Rattus norvegicus) or the Human transcriptomic reference (homo sapiens). The x-axis indicates the percentage of k-mers that are unique to Rat, Human, Both, Neither, or are Ambiguous. More samples are present in the Armenville study than previous analyses because we did not restrict to HMC3-lines.

To further investigate, we traced the cell line’s reported origin in the Baek et al. study. While the paper states that the HMC3 cells were purchased from the Korean Cell Line Bank, the organization confirmed in direct communication that they do not offer this cell line. When contacted, the Baek et al. authors instead referred us to the ATCC website, as reported on their GEO data upload. Given the sourcing ambiguity and shared k-mer content, it is possible that the Baek et al. samples were not fully representative of canonical HMC3 cells or contaminated in some way, which may explain their divergent gene expression and classifier performance.

## Discussion

The HMC3 cell line was created from an in vitro microglial sample in 1995 by transfection of the SV40-T antigen in primary human microglial cultures taken from human embryos (Janabi et al. 1995). At the time of its creation, however, the line was never directly compared to mature human microglial cells to confirm its identity. Complicating its provenance further, HMC3 is reported to be a direct derivative of the CHME-5 cell line, which was later found to be of rat origin (Garcia-Mesa et al. 2017). HMC3 cells were also checked by qualitative immunostaining for the expression of certain markers. Although the HMC3 cells lacked glial fibrillary acidic protein (GFAP) staining and neurofilament staining (used to rule out astrocytic and neuronal contamination), this is not definitive, as other commonly used astrocyte-like lines, such as U87 cells, also lack GFAP expression (Dello Russo et al. 2018). HMC3 also revealed markedly higher basal secretion levels than other microglial cell lines. Additionally, certain chemical transformations done to acquire the immortalized HMC3 cell made it nearly impossible to compare directly with microglial cells (Dello Russo et al. 2018). As HMC3 becomes more widely used, its unexpected behavior has prompted researchers to speculate about its true nature as a microglial substitute. This study’s goal was to provide an informed statistical opinion on the true lineage of the HMC3 cell line.

The results of our study indicate that HMC3 cells are more closely aligned with astrocytic expression profiles than with microglia. Our results demonstrate that the classification method is robust and accurate in our test (Figure 2) and validation sets (Figure 3). Furthermore, we highlight the strength of rules-based features as a viable method to combine gene expression analyses from multiple analyses, i.e., gene-pair qualifiers instead of absolute transcripts per million. While it was thrilling to apply this feature-based tool to multiple datasets with minimum effort in batch correction, the results of this data integration highlight the fact that HMC3 cells classify better with astrocytes than myeloid lineages. One possible explanation is that in the process of creating HMC3, due to the nature of how cells are changed to become immortalized, the cells acquired a stem-like state void of their developmental lineage (Pauklin and Vallier 2013). However, our transformed cell line-based RF classifier refuted this possibility. Therefore, a more likely explanation is that the cell culture from which the HMC3 cell line was created was not pure microglia, and that a contaminating astrocyte gave rise to the SV40-transformed clonal cell line that became known as HMC3. Regardless of the true explanation, our results show that using HMC3 cells as a method to model microglial activity for AD research is not justified.

Our findings align with recent work by Woolf et al. (2025), who conducted a direct phenotypic and functional comparison of commonly used in vitro microglia models, including HMC3 (Woolf et al. 2025). In their study, HMC3 cells failed to express canonical microglia markers (Iba1, CD45, PU.1) and instead stained positive for mural cell markers such as PDGFRβ and NG2. Functionally, HMC3 displayed significantly lower phagocytic activity and secretory responsiveness compared to primary and iPSC-derived microglia—more closely resembling pericytes than cells of myeloid lineage. These findings, derived from an independently sourced and ATCC-validated vial of HMC3 cells, support our transcriptomic results and add further weight to the conclusion that HMC3 is not a valid model of human microglia.

It is critical to accurately determine the origin of HMC3 cell lines, as they are widely used in key research on Alzheimer’s disease (Dello Russo et al., 2018). Microglial recruitment in the brain and central nervous system plays a vital role in supporting brain health and resilience in the face of neurodegenerative diseases, due to microglia’s regulation of several essential immune functions (Miao et al. 2023). This motivated our investigation into the HMC3 line, which we found cannot be accurately classified as microglia. As a result, findings from studies that substitute HMC3 for microglia may be compromised or misleading.

Future studies with larger datasets could provide greater statistical power to detect cell-type-specific expression differences, strengthening confidence in our conclusions. Further analysis should identify the genes that most effectively distinguish astrocytes and microglia from other cell types, and evaluate whether HMC3 has any valid, limited applications. However, our computational findings strongly indicate that HMC3 is not a true human microglial model of myeloid origin, and calls into question its use in prior research and urge caution in continuing to use HMC3 as a proxy for microglia.

## Acknowledgments

The authors would like to thank the countless and often nameless individuals who build, maintain, and provide their sequencing data to the research community. We would also like to thank Samuel Payne, Ph.D., for the tremendous time and effort he spends working with our undergraduate authors and other bioinformatics majors on their capstone projects at Brigham Young University. Brigham Young University’s Office of Research Computing, which provided supercomputing resources for our analysis.

## Scope statement

This study re-evaluates the cellular identity of the HMC3 cell line, which is widely used as a model for human microglia in neurodegenerative disease research. Given previous concerns about its lineage, we developed a gene-pair-based Random Forest classifier using the multiclassPairs R package to analyze bulk and single-cell RNA-sequencing data from multiple publicly available datasets. Our model, trained on over 250 brain-derived samples, demonstrated high classification accuracy and robustness capable of distinguishing among major brain cell types. When applied to HMC3, our analysis consistently predicted a stronger similarity to astrocytes than microglia, challenging its use as a microglial proxy. In contrast, iPSC-derived microglia-like (iMG) cells aligned more closely with primary human microglia, validating our model’s discriminatory power. These findings challenge the validity of HMC3 as a microglial model and highlight the potential for misinterpretation in prior studies. Given the widespread use of HMC3 and its implications for experimental design in investigating disease, we believe these findings warrant immediate dissemination through preprint publication. This work will be of particular interest to researchers studying brain cell identity, RNAseq-based classification methods, and experimental model selection in neurodegenerative disease research.

## Supplementary Figure Captions

**Supplementary Figure 1: multiclassPairs primary tissue classification on training data. (A)** Three subpanels display the results of the classifier on transcription levels from pairs of genes. The top panel contains three rows: “Ref. labels” display cell type assigned by each respective study sample, “Predictions” display the calls of our classifiers on the training set, “Platform/Study” displays how different studies span different cell types. The middle panel displays each sample’s rule activation score (columns). The bottom panel displays the binary heatmap of 452 gene-pair decision rules and the presence and absence of that rule in each sample, black and white, respectively. **(B)** A proximity matrix heatmap portrays similarity clusters of the RF classifier using out-of-bag samples. The top, “Reference labels” display cell type assigned by each respective study, and the “Platform/Study” row displays how different samples span different studies by primary cell types. The heatmap shows pairwise comparisons of the samples’ similarity according to the 452 gene-pair rules, i.e., each sample is given a proximity value with every other sample.

**Supplementary Figure 2: multiclassPairs cell-line classification on training and test data. (A)** Three subpanels display the results of the classifier on transcription levels from pairs of genes. The top panel contains three rows: “Ref. labels” displays the cell-of-origin for the cell type, “Predictions” displays the tool’s classification of the training set, “Sex” displays the cell type’s sex. The middle panel displays each sample’s rule activation score (columns are cell types in DepMap). The bottom panel displays the binary heatmap of 41 gene-pair decision rules and the presence and absence of that rule in each sample, black and white, respectively. **(B)** A proximity matrix heatmap portrays similarity clusters of the RF classifier using in-bag training samples. **(C)** Reflects the sample sub-panel structure as panel A, but for out-of-bag (held-out) testing data. **(D)** Same as panel B, but for the testing data. Only two samples were misclassified, but each has been shown to have a controversial cell of origin in the literature.

