## Supplementary figures and images for "Cellf-Deception: Human microglia clone 3 (HMC3) cells exhibit more astrocyte-like than microglia-like gene expression"

### Supplementary Figures

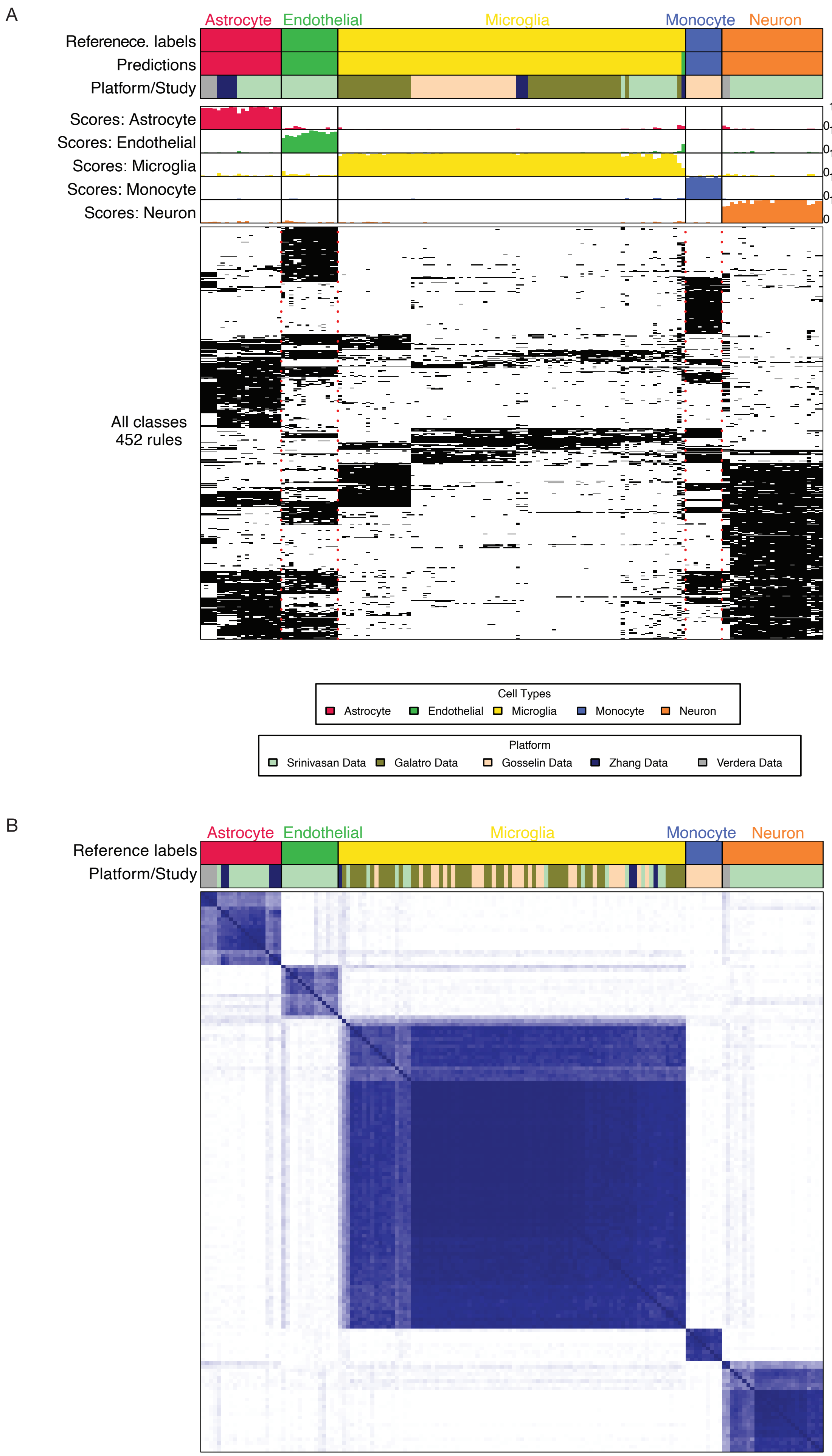

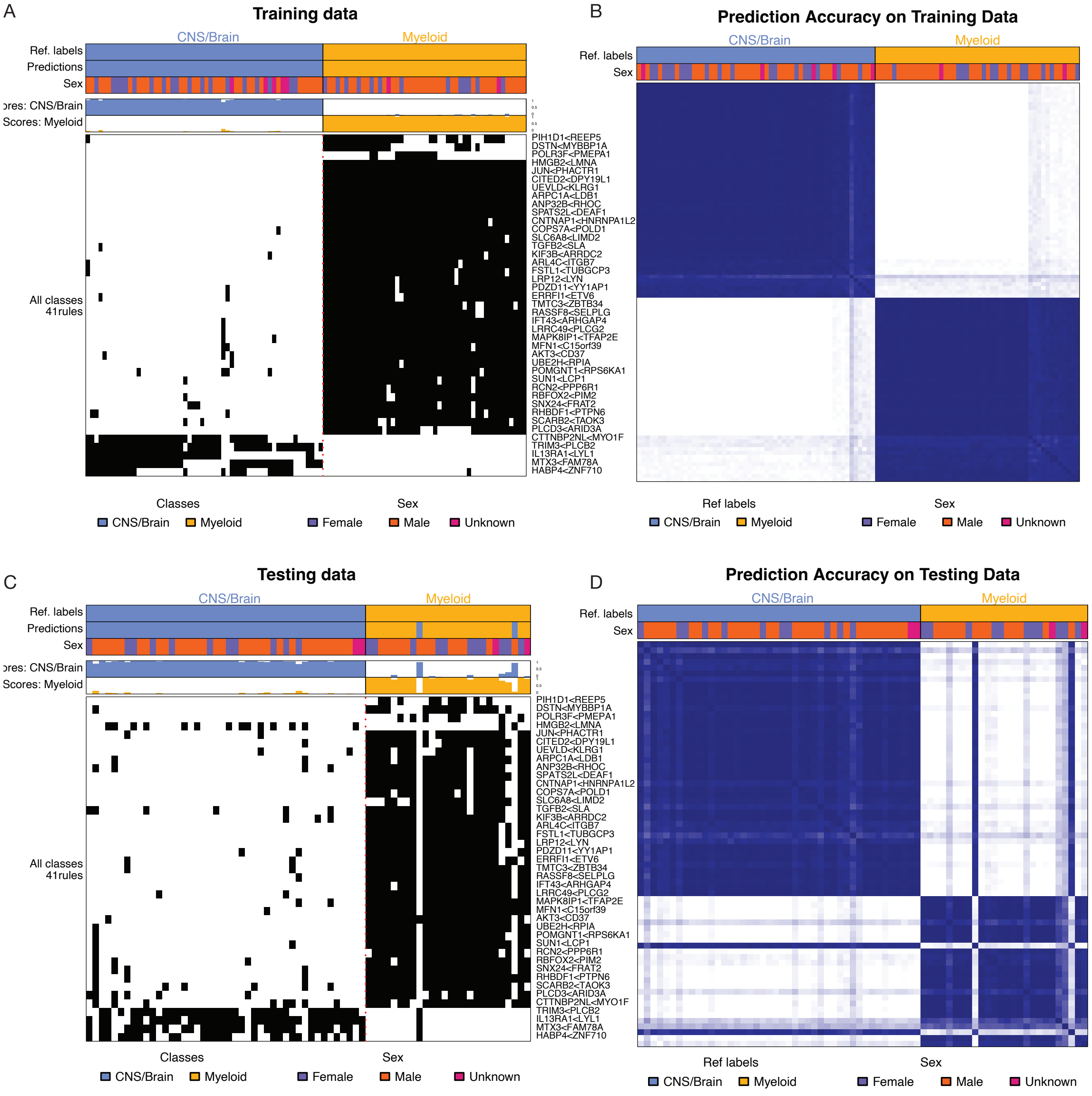
